# Functional coupling between target selection and acquisition in the superior colliculus

**DOI:** 10.1101/2021.04.25.441374

**Authors:** Jaclyn Essig, Gidon Felsen

**Author notes:** Corresponding author: Gidon Felsen.

## Abstract

To survive in unpredictable environments, animals must continuously evaluate their surroundings for behavioral targets, such as food and shelter, and direct their movements to acquire those targets. Although the ability to accurately select and acquire spatial targets depends on a shared network of brain regions, how these processes are linked by neural circuits remains unknown. The superior colliculus (SC) mediates the selection of spatial targets and remains active during orienting movements to acquire targets, which suggests the underexamined possibility that common SC circuits underlie both selection and acquisition processes. Here, we test the hypothesis that SC functional circuitry couples target selection and acquisition using a *default motor plan* generated by selection-related neuronal activity. Single-unit recordings from intermediate and deep layer SC neurons in male mice performing a spatial choice task demonstrated that choice-predictive neurons, including optogenetically identified GABAergic SC neurons whose activity was causally related to target selection, exhibit increased activity during movement to the target. By strategically recording from both rostral and caudal SC neurons, we also revealed an overall caudal-to-rostral shift in activity as targets were acquired. Finally, we used an attractor model to examine how target selection activity in the SC could generate a rostral shift in activity during target acquisition using only intrinsic SC circuitry. Overall, our results suggest a functional coupling between SC circuits that underlie target selection and acquisition, elucidating a key mechanism for goal-directed behavior.

**Significance Statement:** The ability to quickly select and acquire spatial targets is essential to animal survival. Neural circuits underlying these processes are shared in an interconnected network of brain regions, however it is unclear how circuits link decision-making processes with motor commands to execute choices. Here, we examine single-unit activity in the superior colliculus (SC) as mice select and acquire spatial targets to test the hypothesis that choice-related activity promotes target acquisition by generating a default motor plan for orienting movements. By demonstrating that choice-predictive neurons increase their firing rates during movement and capturing the dynamics of SC activity with an attractor model of intrinsic SC circuitry, our results support a role for SC circuits in coupling target selection and acquisition.

## Introduction

Animals continuously evaluate and approach spatial goals while interacting with their environment (Cisek and Kalaska, 2010). Effective goal-directed behavior requires the coordination of sensory, cognitive, and motor processes to accurately identify, select, and acquire spatial targets such as food or mates. Despite being traditionally conceptualized and examined as independent sequential processes whereby decisions are deliberated before actions are planned (Miller *et al.*, 1960), target selection and acquisition in natural environments are fluid processes that continuously influence each other (Cisek and Pastor-Bernier, 2014). However, little is known about how the neural circuits underlying target selection and acquisition are integrated to coordinate goal-directed behavior.

A key node in the network of brain regions mediating goal-directed behaviors is the superior colliculus (SC), an evolutionarily conserved midbrain structure with sensorimotor properties well suited to support target selection and acquisition (Krauzlis *et al.*, 2004; Stein and Stanford, 2008; Gandhi and Katnani, 2011; Wolf *et al.*, 2015; Basso and May, 2017). The intermediate and deep layers of the SC are topographically organized along the rostrocaudal axis such that targets eccentric from the midline are encoded caudally and, as the target is acquired, activity emerges rostrally (Munoz *et al.*, 1991; Anderson *et al.*, 1998; Port *et al.*, 2000; Choi and Guitton, 2009; Gandhi and Katnani, 2011). In general, rostral activity correlates with several aspects of target acquisition, including movement cessation and, in the case of eye movements, microsaccades and fixation (Munoz and Wurtz, 1993; Munoz and Istvan, 1998; Basso *et al.*, 2000; Krauzlis, 2003; Choi and Guitton, 2006; Goffart *et al.*, 2012). Extensive sets of studies have focused on the role of the SC in both target selection (Sparks and Mays, 1980; Horwitz and Newsome, 1999; Glimcher, 2001; McPeek and Keller, 2002; Krauzlis *et al.*, 2004; Felsen and Mainen, 2008; Duan *et al.*, 2015; Basso and May, 2017) and acquisition (Guitton *et al.*, 2003; Gandhi and Katnani, 2011; Marino *et al.*, 2012; Smalianchuk *et al.*, 2018), but it is unclear how these processes are related at the level of SC circuits. One possibility is that distinct sources of cortical and subcortical input to specific cell types along the rostrocaudal axis (Benavidez *et al.*, 2020; Doykos *et al.*, 2020) can mediate selection and acquisition via independent collicular pathways. Another underexamined possibility is that target selection and acquisition are intrinsically linked via SC circuitry. Given the tight linkage between these processes – acquisition must often follow soon after selection – shared neural circuitry could provide behavioral and computational advantages (Wolpert and Landy, 2012; Cisek, 2019). Although much remains unknown about the functional organization of SC circuits, several recent studies have characterized the roles of specific SC cell types in behavior (Masullo *et al.*, 2019; Essig *et al.*, 2020; Wang *et al.*, 2020) and the sophistication of SC circuitry has been increasingly appreciated.

Here, we test the hypothesis that SC circuits are configured such that neural activity underlying the selection of a target promotes a default motor plan for acquiring the target via an orienting movement. Under this hypothesis, we would expect that SC activity causal for target selection would initialize activity for target acquisition which may subsequently be modulated by external input. We recorded rostral and caudal SC activity, including from optogenetically-identified GABAergic neurons, in male mice selecting and acquiring spatial targets via orienting movements. We found that rostral SC neurons reached their peak firing rates during target acquisition, consistent with previous work (Munoz and Wurtz, 1995b; Bergeron *et al.*, 2003; Choi and Guitton, 2009), while many caudal SC neurons exhibited activity related to target selection and remained active during target acquisition. Interestingly, the timing of activity of GABAergic neurons was less dependent on their rostrocaudal location. We next developed an attractor model to examine how intrinsic SC circuitry could promote a shift of activity from caudal to rostral SC during target acquisition. Overall, our findings suggest that SC circuitry may facilitate goal-directed behavior by coupling target selection and acquisition.

## Materials and Methods

Data were collected during experiments previously described in a published study focusing on spatial choice (Essig *et al.*, 2020). All analyses and results presented in the current study are novel. All methodology relevant to the current study is provided here.

### Experimental Design

#### Animals

All procedures were approved by University of Colorado School of Medicine Institutional Animal Care and Use Committee. Mice were bred in the animal facilities of the University of Colorado Anschutz Medical Campus. Heterozygous Gad2-ires-Cre (Gad2-Cre; Gad2^tm2(cre)Zjh/J)^) male mice (n = 10; 6-12 months old at time of experiments) were used in this study. Mice were housed individually in an environmentally controlled room, kept on a 12-hour light/dark cycle and had ad libitum access to food. Mice were water restricted to 1 ml/day and maintained at 80% of their adult weight.

#### Behavioral task

Mice were trained on a previously published odor-guided spatial-choice task (Uchida and Mainen, 2003; Stubblefield *et al.*, 2013; Essig *et al.*, 2020). Briefly, water-restricted mice self-initiated each trial by nose poking into a central port. After a short delay (~200 ms), a binary odor mixture was delivered. Mice were required to wait 500 ± 55 ms (mean ± SD) for a go signal (a high frequency tone) before exiting the odor port and orienting toward the left or right reward port for water (Fig. 1A). We refer to the time between odor valve open and the go signal as the “choice epoch” (Figs. 1D,E; 3). Exiting the odor port prior to the go signal resulted in the unavailability of reward on that trial, although we still analyzed these trials if a reward port was selected. All training and experimental behavioral sessions were conducted during the light cycle.

**Figure 1:**
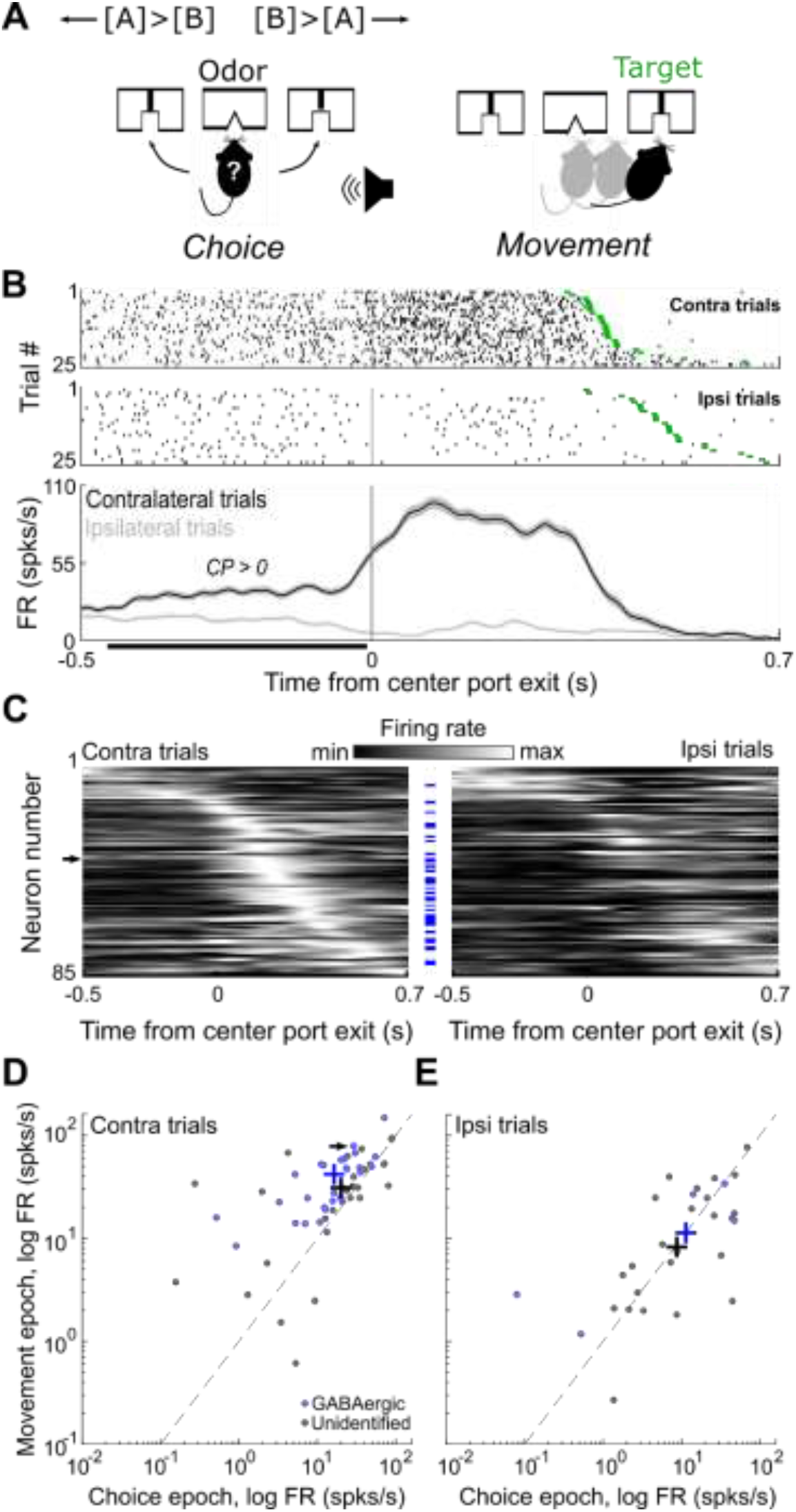
Choice-predictive SC neurons remain active during movement. (**A**) Schematic of task. Mice choose the left or right target (reward port) based on an odor mixture (“Choice”). After a go signal, mice move to the chosen target for a water reward (“Movement”). (**B**) Rasters (top) and peri-event histograms (bottom) for a GABAergic SC neuron that predicts contralateral choice. Black bar indicates the “choice epoch” when the choice prediction of each neuron is calculated based on the area under the ROC curve constructed from the distributions of firing rates on ipsilateral and contralateral trials.(**C**) Normalized activity during choice and movement of choice-predictive SC neurons (n = 85), separately for trials in which the contralateral and ipsilateral target was selected. Neurons are sorted by time of maximum firing rate. Blue dashes indicate GABAergic neurons. Arrow, example neuron in **B**. (**D**) Mean firing rates during the choice epoch plotted against mean firing rates during the movement epoch for each neuron that predicted contralateral choice (n = 55). Only trials in which the contralateral target was selected are shown. Contralateral choice-predictive neurons, including GABAergic neurons, exhibited higher mean activity during movement than choice (*p* < 0.0001, Wilcoxon signed-rank tests). Crosshairs indicate population medians for all (black) and GABAergic (blue) neurons. Arrow, example neuron in **B**. (**E**) As in **D**, epoch for each neuron that predicted ipsilateral choice (n = 30). Only trials in which the ipsilateral target was selected are shown. Activity in this population did not differ between the choice and movement epochs (*p* > 0.05, Wilcoxon signed-rank test).

Odors were comprised of binary mixtures of (+)-carvone (Odor A) and (−)-carvone (Odor B) (Acros), commonly perceived as caraway and spearmint, respectively. In all sessions – including training on the task, as well as during neural recording and manipulation – mixtures in which Odor A > Odor B indicated reward availability at the left port, and Odor B > Odor A indicated reward availability at the right port (Fig. 1A). When Odor A = Odor B, the probability of reward at the left and right ports, independently, was 0.5. The full set of Odor A/Odor B mixtures used was 95/5, 80/20, 60/40, 50/50, 40/60, 20/80, 5/95 and mouse performance on the task was dependent on odor concentration (see Essig *et al*., 2020). Mice completed training in 8-12 weeks and were then implanted with a neural recording drive. All neural recordings were performed in mice that were well-trained on the task.

#### Stereotactic surgeries

Mice were removed from water restriction and had ad libitum access to water for at least one week before surgery. Preparation for surgery was similar for viral injections and chronic implants. Deep anesthesia was induced with 2% isoflurane (Priamal Healthcare Limited) in a ventilated chamber before being transferred to a stereotaxic frame fitted with a heating pad to maintain body temperature. A nose cone attachment continuously delivered 1.3%– 1.6% isoflurane to maintain anesthesia throughout the surgery. Scalp fur was removed using an electric razor and ophthalmic ointment was applied to the eyes. The scalp was cleaned with betadine (Purdue Products) and 70% ethanol before injecting a bolus of topical anesthetic (150 μl 2% lidocaine; Aspen Veterinary Resources) under the scalp. The skull was exposed with a single incision and scalp retraction. The surface of the skull was cleaned with saline and the head was adjusted to ensure lambda was level with bregma (within 150 μm). Immediately following all surgeries, mice were intraperitoneally administered sterile 0.9% saline for rehydration and an analgesic (5 mg/kg Ketofen; Zoetis). A topical antibiotic was applied to the site of incision and mice were given oxygen while waking from anesthesia. Post-operative care, including analgesic and antibiotic administration, continued for up to 5 days after surgery and mice were closely monitored for signs of distress. Additionally, mice recovered after surgery with ad libitum access to water for at least 1 week before being water-restricted for experiments.

For viral injections, a small craniotomy was drilled above the left SC at rostral (3.16-3.4 mm posterior to bregma, 0.75 - 1 mm lateral of midline, 1.8 - 2.17 mm dorsal from the brain surface; n = 3 mice) (Isa *et al.*, 2020) or caudal (3.5 - 4 mm posterior to bregma, 0.8 - 1.25 mm lateral of midline, 0.85 – 1.5 mm dorsal from the brain surface; n = 7 mice) locations (Franklin and Paxinos, 2008). To maximize overlap between ChR2 expression and the optetrode in Gad2-Cre mice, up to 3 injections were made within 0.2 mm^2^. Viruses were delivered with a thin glass pipette at an approximate rate of 100–200 nl/min via manual pressure applied to a 30 ml syringe. Pipets remained at depth for 10 min following each injection before retraction from the brain. Mice were injected with a total (across all injections) of 200 nl of DIO-ChR2-eYFP (AAV2.Ef1α.DIO.ChR2.eYFP, UNC Vector Core, 4.2×10^12^ ppm). After injection, the skin was sutured and mice recovered for 1 week before being water restricted for behavioral training. Expression occurred during the ~10 weeks of training.

To extracellularly record from optogenetically-identified GABAergic SC neurons, an optetrode drive, an optic fiber surrounded by four tetrodes (Anikeeva *et al.*, 2012), was chronically implanted above the DIO-ChR2-eYFP injection site in fully-trained Gad2-Cre mice. A large (~1 mm^2^) craniotomy was made around the initial injection site. Three additional small craniotomies were made anterior to the initial injection site: one for implanting a ground wire and two for skull screws. The drive was slowly lowered into the large craniotomy and secured in place with luting and dental acrylic.

#### Electrophysiology

Extracellular neuronal recordings were collected using four tetrodes (a single tetrode consisted of four polyimide-coated nichrome wires (Sandvik; single-wire diameter 12.5 μm) gold plated to 0.25-0.3 MΩ impedance). Electrical signals were amplified and recorded using the Digital Lynx S multichannel acquisition system (Neuralynx) in conjunction with Cheetah data acquisition software (Neuralynx).

To sample independent populations of neurons, the tetrodes were advanced between 6 - 23 h before each recording session. To estimate tetrode depths during each session we calculated distance traveled with respect to rotation fraction of the thumb screw of the optetrode drive. One full rotation moved the tetrodes ~ 450 μm and tetrodes were moved ~100 μm between sessions. The final tetrode location was confirmed through histological assessment.

Offline spike sorting and cluster quality analysis was performed using MClust software (MClust 4.4.07, A.D. Redish) in MATLAB (2015a). Briefly, for each tetrode, single units were isolated by manual cluster identification based on spike features derived from sampled waveforms. Identification of single units through examination of spikes in high-dimensional feature space allowed us to refine the delimitation of identified clusters by examining all possible two-dimensional combinations of selected spike features. We used standard spike features for single unit extraction: peak amplitude, energy (square root of the sum of squares of each point in the waveform, divided by the number of samples in the waveform), valley amplitude and time. Spike features were derived separately for individual leads. To assess the quality of identified clusters we calculated isolation distance, a standard quantitative metric (Schmitzer-Torbert *et al.*, 2005). Clusters with an isolation distance > 6 were deemed single units. Units were clustered blind to interspike interval, and only clusters with few interspike intervals < 1 ms were considered for further examination. Furthermore, we excluded the possibility of double counting neurons by ensuring that the waveforms and response properties sufficiently changed across sessions. If they did not, we conservatively assumed that we recorded twice from the same neuron, and only included data from one session.

Electrophysiological recordings were obtained from 285 SC neurons in 96 behavioral sessions. Details of our analyses of the data obtained from our recording experiments are described below. All neural data analyses were performed in MATLAB (2015a/2019a). Neurons recorded during behavior/recording sessions with fewer than 40 trials in either direction or with a choice-epoch and movement-epoch firing rate below 2.5 spikes/s for trials in both directions were excluded from all analyses, resulting in 285 neurons in this data set.

#### Histology

Final tetrode location was confirmed histologically (see Essig *et al.*, 2020). Mice were overdosed with an intraperitoneal injection of sodium pentobarbital (100 mg/kg; Sigma Life Science) and transcardially perfused with phosphate buffered saline (PBS) and 4% paraformaldehyde (PFA) in 0.1 M phosphate buffer (PB). After perfusion, brains were submerged in 4% PFA in 0.1 M PB for 24hr for post-fixation and then cryoprotected for at least 12hr immersion in 30% sucrose in 0.1 M PB. On a freezing microtome, the brain was frozen rapidly with dry ice and embedded in 30% sucrose. Serial coronal sections (50 μm) were cut and stored in 0.1M PBS. Sections were stained with 435/455 blue fluorescent Nissl (1:200, NeuroTrace; Invitrogen) to identify cytoarchitectural features of the SC and verify tetrode tracks to determine the depth and rostrocaudal location of the recordings.

#### Optogenetic identification of GABAergic neurons

Before and/or after behavioral sessions, light was delivered via a diode-pumped, solid-state laser (473 nm; Shanghai Laser & Optics Century) at 8 Hz (10 ms on) or, for a small sample (< 20 sessions), at 25 Hz (10 ms on) to record extracellular light-induced activity from the same population of neurons that were recorded during behavior. Isolated units were identified as GABAergic based on reliable, short-latency responses to light and high waveform correlations between spontaneous and light-evoked action potentials (see Essig *et al*., 2020 for further details). Neurons identified as GABAergic could be further identified (i.e., “tracked”) during behavioral sessions based on spike features and location (i.e., tetrode number and lead number).

#### Choice prediction

To examine the dependence of the firing rate of individual neurons on choice (Fig. 1C-E), we used an ROC-based analysis (Green and Swets, 1966) that quantifies the ability of an ideal observer to classify whether a given spike rate during the choice epoch was recorded in one of two conditions (here, during ipsilateral or contralateral trials)(Feierstein *et al.*, 2006; Essig *et al.*, 2020). We defined the choice epoch as beginning 100 ms after odor valve open and ending with the go signal. Statistical significance was determined with a permutation test: we recalculated choice prediction (CP) after randomly reassigning all firing rates to either of the two groups arbitrarily, repeated this procedure 500 times to obtain a distribution of values, and calculated the fraction of random values exceeding the actual value. We tested for significance at α = 0.05. Using this analysis we found that 55/285 neurons predicted contralateral choice (CP > 0, *p* < 0.05) and 30/285 neurons predicted ipsilateral choice (CP < 0, *p* < 0.05) (Fig. 1C-E); see Essig *et al.* (2020) for more details).

#### Comparing activity during choice and movement epochs

To examine activity across the choice and movement epochs of choice-predictive neurons (*p* < 0.05), activity was averaged across contralateral and ipsilateral trials (separately) in 10 ms bins (Fig. 1C). The activity of each neuron was rescaled across both directions and epochs from 0 (lowest activity; black) to 1 (highest activity; white), smoothed with a gaussian filter (σ = 40 ms) and sorted by timing of maximum firing rate. Example peristimulus time histograms in Figure 1B were smoothed with a Gaussian filter (σ = 10 ms).

We compared median population firing rates for all choice-predictive neurons and GABAergic choice-predictive neurons between the choice epoch (100 ms after odor delivery to go signal; ~400 ms) and movement epoch (center port exit to target port; ~630 ms) separately for contralateral choice-predictive and ipsilateral choice-predictive neurons (Fig. 1D,E). Neurons were included for analysis if their mean firing rate across the sessions was > 1 spk/s for at least 1 epoch (choice or movement) in the direction analyzed (ipsilateral or contralateral trials). Mann-Whitney U tests were performed for all comparisons.

To examine firing rate throughout the movement epoch, activity was averaged across contralateral and ipsilateral trials (separately) in 10 ms bins (Fig. 2A). The activity of each neuron was rescaled across both directions and both epochs from 0 (lowest activity; black) to 1 (highest activity; white), smoothed with a gaussian filter (σ = 20 ms) and sorted by the time of maximum firing rate. Neurons with peak firing rates at the time of target entry were omitted from analysis (n = 42 neurons).

**Figure 2:**
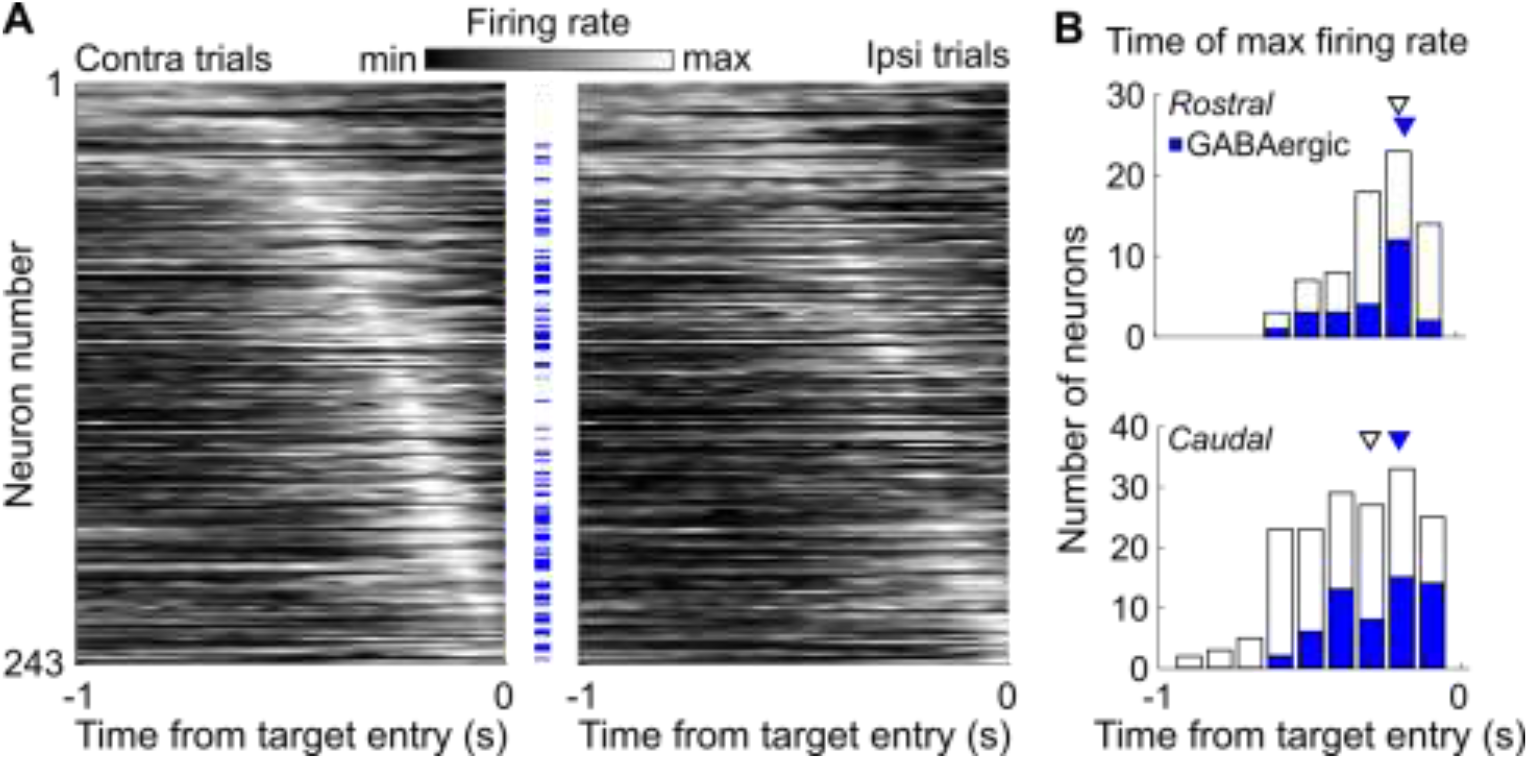
Timing of SC activity during movement to the target. **(A)** Normalized activity of all SC neurons during movement, separately for trials in which the contralateral and ipsilateral target was selected. Neurons are sorted by time of maximum firing rate and omitted from analysis if their peak firing rate occurred at the time of target entry (t = 0 s). Blue dashes indicate GABAergic neurons. Peak firing rate of GABAergic neurons occurs later than the overall population (*p* = 0.021, Mann-Whitney U test). (**B**) Time of maximum firing rate separately for rostral and caudal populations. In the overall population, peak firing rate was later in rostral (top) than caudal neurons (bottom) (*p* = 0.00066, Mann-Whitney U test). Arrowheads show median time of peak firing rate for overall (open) and GABAergic (blue) populations of neurons.

#### Attractor model

We adapted a rate-based bump attractor model originally developed to study spatial choice (Lintz *et al.*, 2019; Essig *et al.*, 2020) to examine rostrocaudal activity dynamics during movement. As detailed in Essig *et al.* (2020) and briefly described here, the model consists of 200 excitatory (*E*) cells and 100 inhibitory (*I*) cells per SC (600 cells total). Intra-SC synaptic weights were larger for nearby cells, and smaller for more distant ones, determined by

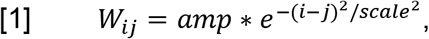

where *i* and *j* are the locations of the pre- and post-synaptic cells, respectively, and *amp* and *scale* are defined independently for presynaptic *E* and *I* cells (*ampE* = 0.01, *ampI* = 0.15, *scaleE* = 0.03). *amp* sets the amplitude (i.e., “strength”) of the connection weights, and *scale* determines the spatial extent over which the connection strength decays. Overall, *I* cells had a higher *scale* than *E* cells, with *scaleI* magnitude determined by the rostral (*I* cells 1:25) or caudal (*I* cells 26:100) location of the presynaptic cell: *scaleI_rostral_* = 0.5 and *scaleI_caudal_* = 0.4. Inter-SC synaptic connections were made sparse by assigning synaptic weights to a subset of probabilistically determined contralateral *E* and *I* cells and were reciprocated between the left and right SC. To promote network stability, each *W* was normalized to have a maximum eigenvalue of 1.5 by dividing all connection values by max(λ)/1.5, where max(λ) is the largest eigenvalue of the matrix after initialization.

Spike rates of *E* and *I* cells in the left SC (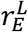 and 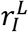, respectively) evolved at each time step (~ 2 ms in our numerical simulations) according to

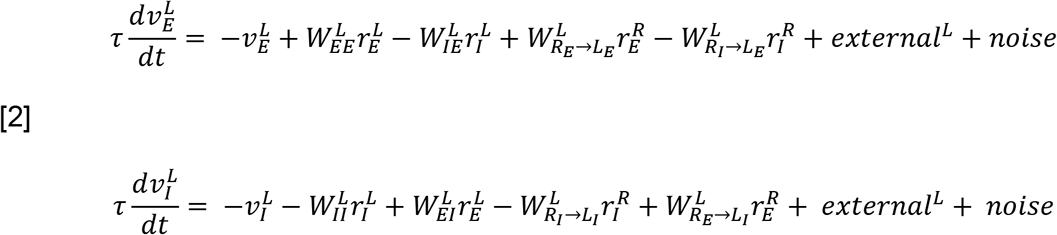

and all spike rates were rectified at each time step according to

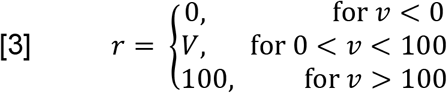

where, e.g., 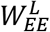 represents synaptic weights from left *E* to *E* cells, 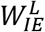 represents weights from left *I* to *E* cells, and 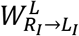 represents weights from right*I* to left *E* cells. Noise was drawn from a Gaussian distribution with mean = 0, and variance = 15 for *E* cells and 5 for *I* cells. 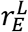 and 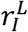 are vectors, with one entry per *E* or *I* cell in the left SC, respectively. Similarly, 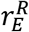 and 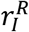 describe the firing rates of cells in the right SC, and they evolve over time via the same equations as those in the left SC (i.e., via Eqs. [2], [3], with all “L”s and “R”s swapped).

The vectors *external*^*L*^ and *external*^*R*^ represent the drive to the *E* and *I* cells from sources outside the SC. For all trial types, external drive was applied in a linearly graded fashion to a small fraction of the most caudal cells (*E* cells numbered 170 to 179 and *I* cells numbered 85 to 89). On leftward trials, the right SC received a stronger external drive than the left SC, and vice versa on rightward trials. For each trial, external drive began at time step 151, corresponding to the odor delivery time in the task, and stopped time step 400 for an approximate total time of ~ 500 ms. At step 401, immediately following removal of the external input, choice was determined based on the SC with the highest average caudal *E* cell firing rate. Each “session” consisted of 255 trials, with the proportion of different difficulties matched to the behavioral sessions. Each model data point consists of 20 sessions (Fig. 4C,D; 5A,C).

**Figure 3:**
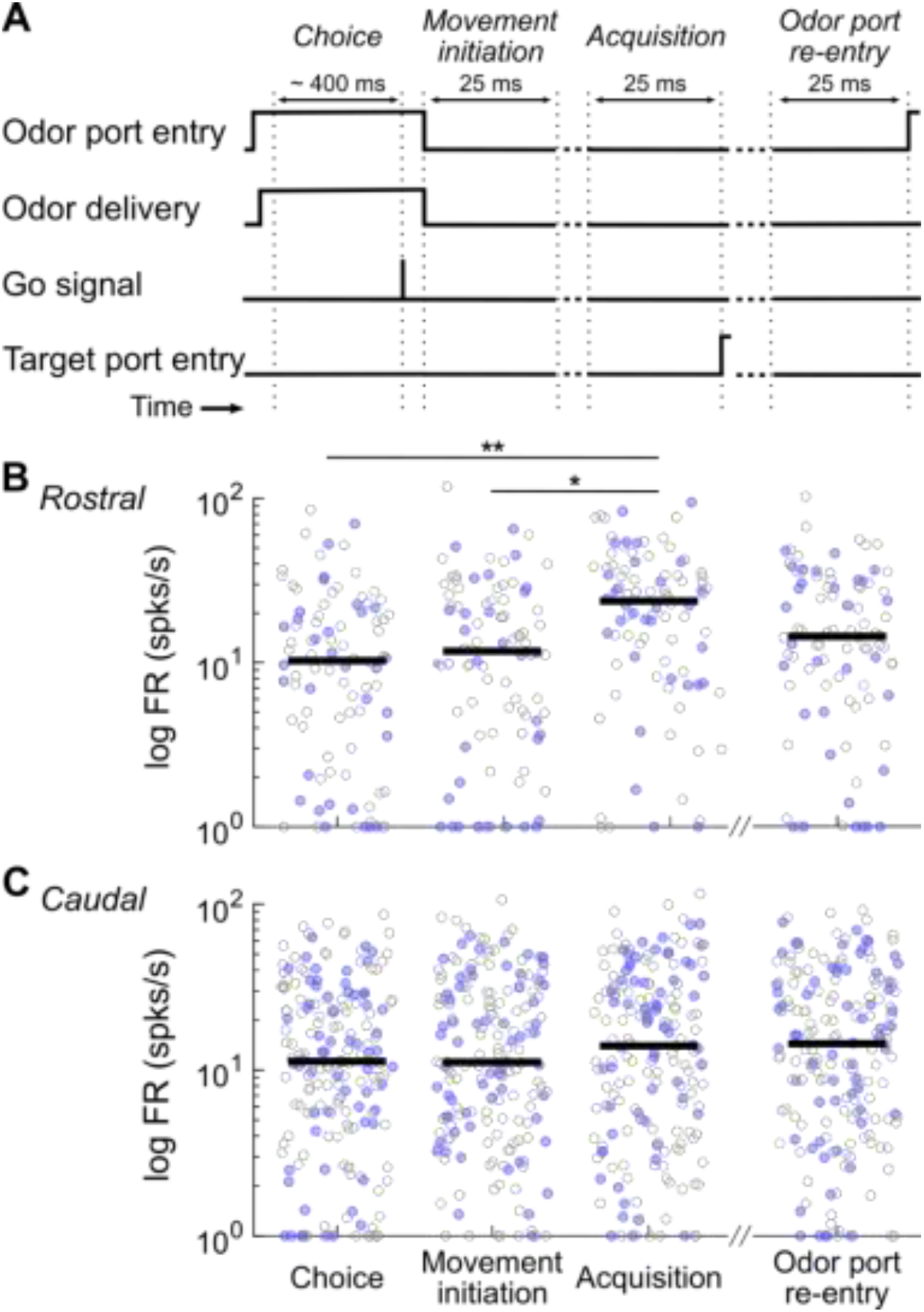
Comparison of SC activity across task epochs. (**A**) Definitions of event-based epochs of interest. (**B**) Mean activity of all rostral neurons on trials in which the contralateral target was selected. Blue circles indicate GABAergic neurons. Black bars, median firing rate of all rostral neurons. ** *p* = 0.0004, * *p* = 0.024, multiple comparisons. Results were similar for trials in which the ipsilateral target was selected (not shown). (**C**) As in **B**, for caudal neurons.

**Figure 4:**
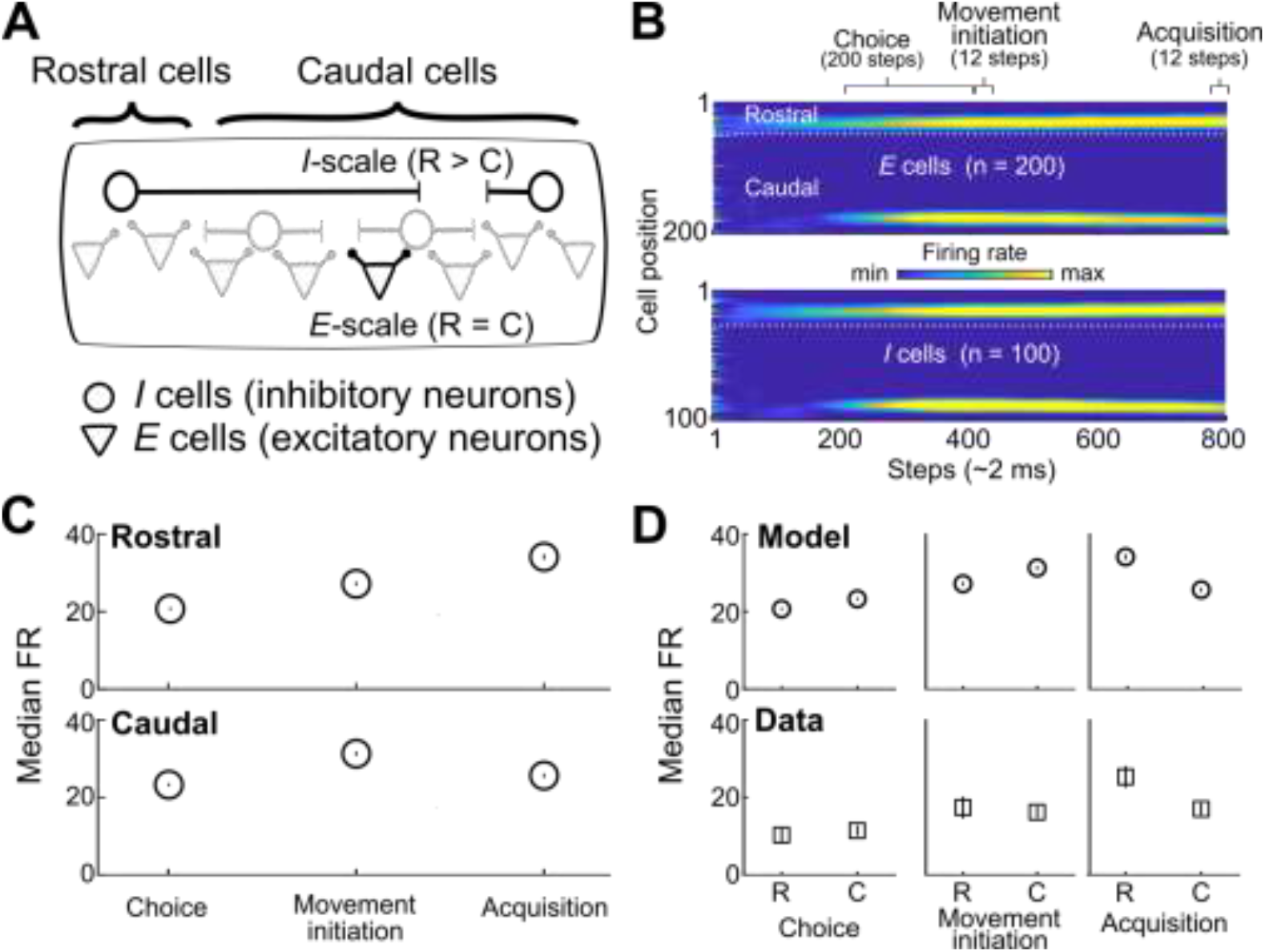
Attractor model captures spatiotemporal dynamics of neural activity during the task. (**A**) Schematic of model. *I-* and *E*-scale refer to range of inhibition and excitation, respectively, over which the strength of the synaptic influence decays. Rostral neurons comprise the first quarter of cells, the rest are caudal. (**B**) Example activity of excitatory (*E*) cells (top) and inhibitory (*I*) cells (bottom) on a single trial. Epochs were identified for comparison with neural activity during the task. External excitation is delivered to the most caudal *E* and *I* cells only during the choice epoch. Dotted line shows division between rostral and caudal populations. (**C**) Median firing rates for each epoch on contralateral trials of rostral (top) and caudal (bottom) *E* cells averaged over 20 sessions. Error bars, +/− SEM. (**D**) Median rostral and caudal firing rates across epochs from the model (top) and from neural recordings (bottom).

**Figure 5:**
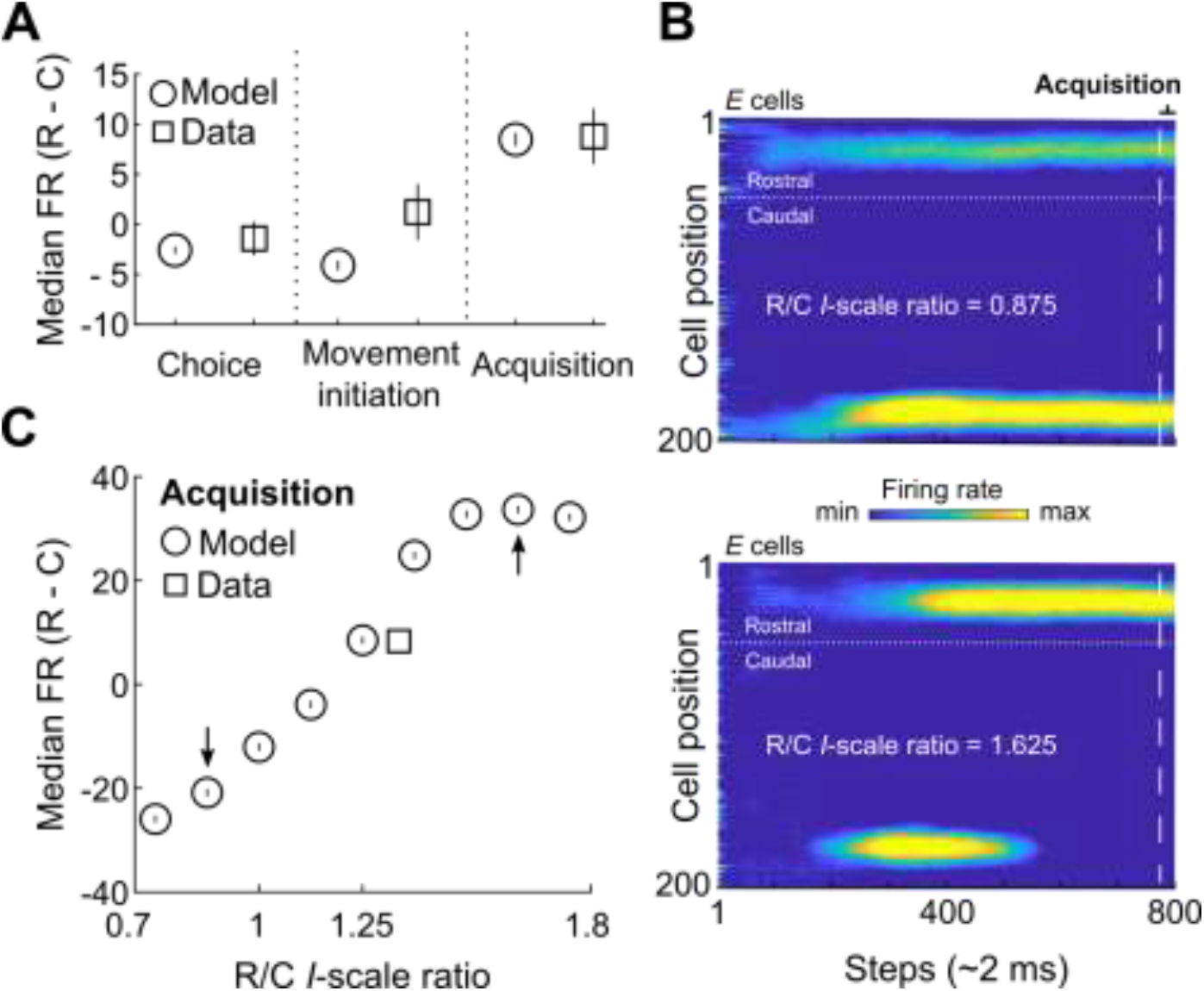
Rostrocaudal activity during target acquisition depends on relative spatial influence of rostral and caudal inhibitory cells. (**A**) Difference between rostral and caudal (R-C) median firing rates for the model and neural recordings for each epoch. (**B**) Example activity of model *E* cells for two R/C *I*-scale ratios on a single trial when caudal *I*-scale is greater than rostral *I*-scale (top) and rostral *I*-scale is far greater than caudal *I*-scale (bottom). Dashed lines indicate beginning of acquisition epoch. (**C**) Difference in rostral and caudal median *E*-cell firing rates during the acquisition epoch for a range of rostral/caudal (R/C) *I*-scale ratios. Data from neural recordings are plotted as a square for comparison. Arrows indicate *I*-scale ratios used in **B.**

For firing rate calculations for E cells (Figs. 4; 5), the duration of each epoch matched epochs in the task and were defined as follows: choice epoch, steps 200-399; movement initiation, steps 400-11; acquisition, steps 789-800.

#### Statistical Analysis

In general, normality of distributions was tested using the Anderson-Darling test. Data were analyzed using MATLAB (2015a/2019a) and presented primarily as medians or means ± SEM. *p* values for each comparison are reported in the figure legends, results and/or methods sections. Mice were excluded from analyses based on the misalignment of the optetrode with ChR2 expression, lack of ChR2 expression, and misplacement of the optetrode outside of the SC (13 mice were excluded from the final data set).

#### Code Accessibility

The data that support the findings of this study and the custom MATLAB code are available from the corresponding author upon written request.

## Results

### Activity in choice-predictive SC neurons increases during movement to selected targets

To begin testing the hypothesis that selecting a target promotes a default motor plan in the superior colliculus (SC) for acquiring the target, we examined how neural activity evolved in the SC of freely-moving mice selecting and acquiring targets in a two-alternative spatial choice task (Stubblefield *et al.*, 2013; Essig *et al.*, 2020). The task required mice to first enter a central port and to sample a binary mixture of odors. Mice were trained to select either the left or right reward port based on the dominant component of the odor mixture (“Choice”; Fig. 1A) and, at the presentation of a go signal, initiate an orienting movement from the center port to the selected reward port to retrieve a water reward (“Movement”; Fig. 1A). As mice performed the task, we obtained single-unit recordings, including from identified GABAergic neurons (Materials and Methods; see Essig *et al*., 2020), from the intermediate and deep layers of the SC. The dissociation of choice and movement epochs enabled us to compare choice- and movement-related activity in our recorded population. Consistent with previous reports, including of primates performing saccadic eye movements (Munoz and Wurtz, 1995a; Basso and Wurtz, 1998; Everling *et al.*, 1999; Felsen and Mainen, 2008), many neurons displayed activity throughout choice and movement (Fig. 1B).

We next examined the movement activity of a subset of choice-predictive neurons that likely causally contribute to target selection (Essig *et al.*, 2020). Briefly, neurons are classified as choice predictive based on an ROC analysis of their firing rates during the choice epoch (Materials and Methods). For example, neurons that exhibit significantly higher firing rate when selecting the contralateral than the ipsilateral target would be classified as contralateral choice-predictive neurons. We found that choice-predictive neurons (n = 85/285 neurons, p < 0.05) appeared to be highly active during movement (Fig. 1C). We quantified the change in activity of these neurons between the choice epoch (during odor sampling; ~ 400 ms) and the movement epoch (from center port to reward port; ~630 ms). Since the activity of many SC neurons depends on movement direction (Horwitz and Newsome, 2001; Felsen and Mainen, 2008; Essig *et al.*, 2020), we examined activity for ipsilateral and contralateral trials separately. We first compared the firing rates for the choice epoch to the movement epoch on contralateral trials and found that for our total population of contralateral choice-predictive neurons, as well as for contralateral choice-predictive neurons identified as GABAergic, activity significantly increased from choice-related levels during movement (p < 0.0001, Wilcoxon signed-rank test; Fig. 1D). We did not find a similar increase in activity during the movement epoch for ipsilateral choice-predictive neurons on ipsilateral trials (*p* >0.05, Wilcoxon signed-rank test; Fig. 1E). These results demonstrate that SC neurons mediating target selection are more active during movement than during choice, even in identified GABAergic populations, consistent with the hypothesis that choice-related activity is linked to a motor plan for target acquisition in the SC.

### Rostral shift of activity during movement

Given the high activity during movement, even among neurons likely involved in target selection (Essig *et al.*, 2020), we next examined how the activity of our entire population of SC neurons evolves as mice are orienting to, and acquiring, spatial targets. We found that the majority of neurons were active during movement and, across the population, peak activity was exhibited throughout the duration of the movement (Fig. 2A). Interestingly, population activity during contralateral and ipsilateral orienting movements was similar, although, again consistent with a prominent role for the SC in mediating movements to contralateral targets (Gandhi and Katnani, 2011), the maximum firing rate for most neurons occurred during contralateral trials (n = 144/243 neurons). The activity of GABAergic neurons (Fig. 2A, blue dashes) tended to peak later than the overall population (total population median: 250 ms before target entry; GABAergic population median: 190 ms; *p* = 0.021, Mann-Whitney U test).

We next analyzed the timing of maximum firing rate based on the rostrocaudal location of each neuron as previous studies have postulated that activity shifts from caudal to rostral SC as targets are acquired (Munoz *et al.*, 1991; Guitton *et al.*, 2003). By targeting our recordings to the rostral or caudal SC in separate sets of mice, we recorded from 73 rostral neurons (25 GABAergic) and 170 caudal neurons (58 GABAergic).

In the overall population, rostral neurons reached their maximum firing rates later than caudal neurons (Fig. 2B, white arrowheads; rostral: 190 ms before target entry; caudal: 245 ms; *p* = 0.00066, Mann-Whitney U test). These results are consistent with SC activity recorded from cats and primates performing similar tasks (Munoz *et al.*, 1991; Munoz and Wurtz, 1995b, 1995a) and suggest that, as a population, SC represents movements to spatial targets. Interestingly, the activity of GABAergic neurons did not depend on rostrocaudal location and were active significantly later during movement than the total population (Fig. 2B, blue arrowheads; rostral: 170 ms before target entry; caudal: 205 ms; *p* = 0.92, Mann-Whitney U test), consistent with a role for GABAergic neurons in target acquisition, in addition to their role in target selection (Essig *et al.*, 2020).

### Rostral SC activity increases as targets are acquired

To more rigorously examine differences in activity between rostral and caudal neurons, we quantified their activity in epochs capturing key features of the task: choice, movement initiation, target acquisition and odor port re-entry (Fig. 3A). As in Figure 1D and E, the choice epoch was defined as when mice are in the center port and sampling the odor (~400 ms; Fig. 3A). The remaining epochs were 25 ms each and yoked to specific task events: “movement initiation” (the 25 ms following odor poke out), “acquisition” (the 25 ms preceding port entry) and “odor port re-entry” (the 25 ms preceding odor port entry; Fig. 3A). We focused on contralateral trials since these generally elicited higher activity than ipsilateral trials (Figs. 1C; 2A). Firing rates for GABAergic neurons did not differ from that of the overall population in any epoch (p > 0.05, Mann-Whitney U tests); we thus performed our inter-epoch comparisons on the overall population.

We first examined the activity of neurons recorded in the rostral SC (Fig. 3B). Overall, we found that rostral activity, while exhibiting significant variability within epochs, was modulated between epochs in the task (Fig. 3B; *p* = 0.00077, Kruskal-Wallis test). Population activity of rostral neurons remained unchanged from choice to movement initiation (*p* = 0.66), however, there was a significant increase in activity during acquisition compared to choice (*p* = 0.00041) and movement initiation (*p* = 0.023, multiple comparison tests). Rostral activity during odor port re-entry did not differ from that during acquisition or the other epochs, perhaps reflecting the fact that, within the context of the task, entering the odor port could be considered a form of target acquisition, but was only indirectly linked to the goal of obtaining water. Conversely, neurons recorded from the caudal SC maintained choice-epoch firing rates for each subsequent epoch (Fig. 3C; *p* = 0.46, Kruskal-Wallis test). Together, these results reveal that activity in caudal SC persists throughout the selection and acquisition of spatial targets whereas rostral activity shows an increase in activity specifically as targets are acquired.

### Intrinsic SC circuitry can support a rostral increase in activity for target acquisition

Our results demonstrate that most choice-predictive neurons exhibit higher activity during movement to the target than during choice (Fig. 1) and that activity in rostrally recorded neurons increases during target acquisition (Figs. 2; 3). If choice-related activity initiates a default motor plan for subsequent acquisition of selected targets, we would expect that intrinsic SC circuitry would suffice to link choice to acquisition without input from extracollicular structures (e.g., from the substantia nigra or cerebellum, both of which are known to modulate movement-related SC activity). To test for network architectures that may link target selection and acquisition in the SC, we extended an attractor model initially developed to interrogate intrinsic SC circuits for decision making (Fig. 4)(Essig *et al.*, 2020). Briefly, the model includes a left and a right SC, each with their own populations of excitatory (*E*) and inhibitory (*I*) cells. Cells are arranged linearly, with rostral cells comprising the first quarter of the population and the remaining cells classified as caudal (Fig. 4A). After external excitation representing the odorants is applied to *E* and *I* cells during the time steps corresponding to the choice epoch, external input is removed and the choice is read out based on the overall activity of the caudal *E* cells on the following step of the model (Essig *et al.*, 2020).

We first extended the model by identifying parameters that would sustain activity well beyond the choice epoch while also preserving choice accuracy. We accomplished this by reducing the synaptic scale of *E* cells such that recurrent excitation occurred over a smaller area and could be maintained for longer durations. However, this change did not produce a rostrally-directed shift in activity during movement, as we observed experimentally (Figs 2; 3). Therefore, guided by previous reports of long-range inhibitory interactions between rostral and caudal populations (Meredith and Ramoa, 1998; Munoz and Istvan, 1998; Behan *et al.*, 2002) and rostrocaudal inhibitory asymmetries in the SC (Bayguinov *et al.*, 2015), we introduced asymmetry to the inhibitory projections based on the rostrocaudal position of the inhibitory cells (Fig. 4A). Interestingly, when we increased the spatial influence of rostral *I* cells (*I* – scale), activity emerged during choice in caudal and rostral populations (Fig. 4B), despite providing external input exclusively to caudal cells during the choice epoch.

To determine how modeled activity evolved from target selection to acquisition, we identified model epochs that were analogous to the epochs used for our experimental data (Fig. 4B). Each step in the model is equivalent to approximately 2 ms of biological time; thus, we could estimate when the movement initiation and target acquisition would occur in relation to the choice epoch and match their durations to the epochs used for the experimental data (Fig. 4B). Similar to our experimental results, rostral activity increased for the acquisition epoch whereas caudal activity remained relatively stable for each epoch following the choice epoch (Fig. 4C). Figure 4D compares the activity of rostral and caudal populations for each epoch in the model (top) and for the experimental data (bottom). In general, the model was able to recapitulate the relative activity across epochs in the rostral and caudal populations, including the increase in rostral activity during target acquisition.

We next examined how the model results depended on the built-in rostrocaudal inhibitory asymmetry (Fig. 4A). Since we were primarily interested in the relative difference in firing rate between rostral and caudal SC populations, we directly compared the model results to the data by calculating the difference in rostral and caudal firing rates for each epoch (Fig. 5A). We then increased or decreased the rostral *I*-scale parameter, re-ran the model, and focused on activity during the acquisition epoch, in which we observed the greatest relative increase in rostral activity (Fig. 5A). Decreasing the rostral *I*-scale parameter resulted in less activity accumulating rostrally during acquisition (Fig. 5B, top) whereas increasing the rostral *I*-scale parameter resulted in a pronounced increase in rostral activity and a severe decrease in caudal activity (Fig. 5B, bottom). We repeated this analysis for a range of values of the rostral *I*-scale parameter to reveal that our results depended strongly on the rostral-caudal *I-*scale ratio (Fig. 5C), suggesting a possible dependence of the rostrocaudal balance of activity on the relative spatial influence of rostral and caudal inhibitory SC cells.

Overall, the model demonstrates that choice-related activity and intrinsic SC circuitry alone, without external modulation, could account for target-acquisition-related activity and further supports the hypothesis that functional SC circuits underlying target selection provide a default motor plan for target acquisition.

## Discussion

In this study, we examined the idea that shared neural circuitry underlies the selection of targets for movement and their acquisition. We hypothesized that, since circuits in the superior colliculus (SC) mediate target selection and acquisition (Mays and Sparks, 1980; Horwitz and Newsome, 1999; Glimcher, 2001; Guitton *et al.*, 2003; Felsen and Mainen, 2008; Gandhi and Katnani, 2011; Duan *et al.*, 2015; Basso and May, 2017), SC circuitry links these processes by establishing a default motor plan for target acquisition initiated by selection-related activity. By recording from rostral and caudal SC neurons while mice selected, oriented to, and acquired targets, we found that many neurons were active for these task-related behaviors, and that activity was generally higher during movement, even in those neurons thought to play a causal role in target selection (Fig. 1). The balance of activity shifted rostrally during movement, although not among the GABAergic population that was more active later in the movement regardless of rostrocaudal position (Fig. 2), and an attractor model limited to intrinsic SC circuits accounted for this rostral shift (Fig. 4). Together, these results are consistent with the idea that functional circuitry for target selection in the SC provides a default motor plan for subsequently acquiring the selected target.

Our work builds on a wealth of studies examining how SC activity relates to target selection and acquisition, which have mainly focused on gaze shifts (Sparks, 1999), primarily via saccadic eye movements, in primates and cats (Gandhi and Katnani, 2011). The full-body orienting movements required by our task appear to similarly engage the mouse SC: Perturbation experiments predictably bias choices (Felsen and Mainen, 2008; Stubblefield *et al.*, 2013; Lintz *et al.*, 2019; Essig *et al.*, 2020), and we observed that activity in rostral SC tended to peak later during movement (Fig. 2B), and was higher during target acquisition (Figs. 3B,C; 4D), than activity in caudal SC. Data from the SC of primates and cats making saccades is broadly similar (Wolf *et al.*, 2015), however some of our results differed from previous data. For example, we found that caudal SC activity was only slightly higher than rostral activity during the choice epoch and was even higher during acquisition than during choice (Fig. 3B,C). These results are inconsistent with a strict caudal-to-rostral shift in the locus of activity during movement, whereby caudal activity decreases as rostral activity increases, as has been described for gaze shifts (Guitton *et al.*, 2003). Instead, our results show that rostral activity is more strongly modulated by target acquisition than caudal activity, driving a rostral spread of activity during movement without a concomitant decrease in caudal activity (Anderson *et al.*, 1998). Thus, while we found that the balance of SC activity shifts rostrally during movement, consistent with previous results in primates and cats during saccades, activity in caudal SC during full-body movements in mice may differ from caudal SC activity in primates and cats during saccades. To examine this possibility further, simultaneous recordings at precise locations along the rostrocaudal axis would need to be performed.

Our study was the first to examine the activity of GABAergic SC neurons during target acquisition. We found that GABAergic neurons are active slightly later during movement than the overall population of SC neurons (Fig. 2), but that their activity during target acquisition was broadly similar to the overall population (Fig. 3B,C). These results suggest that, similar to their role during target selection (Essig *et al.*, 2020), GABAergic neurons do not primarily provide local suppression: If they did, we would expect to observe qualitative differences between the activity of GABAergic and the overall population. Rather, they may provide longer-range inhibition during target acquisition. Consistent with the role of inhibition in our attractor model (Fig. 4) and a proposed mechanism for stopping saccades (Munoz and Istvan, 1998), GABAergic neurons may direct SC activity rostrally as movement occurs to ultimately promote movement cessation at the target.

While our results support the idea that target acquisition is influenced by target selection via shared circuitry in the SC, they do not rule out a role for other brain regions in modulating target acquisition-related activity in the rostral SC. Indeed, the rostral SC integrates input from numerous brain regions (Benavidez *et al.*, 2020; Doykos *et al.*, 2020), many of which could contribute to processes related to target acquisition, including regulating movement speed and movement cessation. In particular, the cerebellar nuclei are thought to play a role in online motor control by modulating activity in movement-related circuitry, suggesting that their robust projections to the rostral SC contribute to target acquisition for SC-dependent orienting movements (Noorani and Carpenter, 2017). This input may be critical for retuning target acquisition when real-time sensorimotor remapping is required or may increase the precision of target acquisition under stable conditions. Thus, while a default motor plan for target acquisition in the SC may be formed during target selection, the circuitry is sufficiently flexible to allow for modulation by other structures.

Our findings raise several questions that can be addressed by future studies. First, while our study and others have shown that activity in rostral SC correlates with target acquisition, it is unclear whether this relationship is causal. Electrical microstimulation and pharmacological inhibition of primate rostral SC during goal-directed eye movements affects the movement endpoint (Gandhi and Keller, 1999; Basso *et al.*, 2000; Hafed *et al.*, 2008; Goffart *et al.*, 2012), but the effect of transiently inhibiting rostral SC activity during movement is not known. Restricting inhibition to specific cell types and during specific phases of movement (e.g., optogenetically) could elucidate the functional neural circuitry in SC necessary for target acquisition. In particular, the GABAergic SC neurons studied here comprise a heterogeneous population (Mize, 1992; Sooksawate *et al.*, 2011) that plays multiple functional roles. Recording and perturbing specific types of GABAergic neurons, as well as other types of SC neurons such as those projecting to brainstem motor nuclei, during target acquisition would be valuable. In addition, our analyses examined how SC circuitry could guide orienting movements to the target regardless of the specific effectors involved, consistent with the framework that the SC performs a general role in representing spatial targets (Krauzlis *et al.*, 2004). Since the SC mediates pinnae movements (Stein and Clamann, 1981), whisking (Hemelt and Keller, 2008), licking (Rossi *et al.*, 2016) and head-orientation (Corneil *et al.*, 2002; Wilson *et al.*, 2018), all of which undoubtedly occur during our task, future studies can use high-resolution tracking methods (Mathis and Mathis, 2020) while recording SC activity to examine how these body adjustments are encoded in SC circuits as animals acquire targets. Finally, the functions of external inputs to the rostral SC during target acquisition can be examined, towards the larger goal of understanding how the SC integrates input from a network of brain regions to coordinate goal-directed behavior.

## Acknowledgements

This work was supported by the National Institutes of Health (R01NS079518, F31NS103305), Rocky Mountain Neurological Disorders Center (P30NS048154), by NIH/NCRR Colorado CTSI grant UL1 RR025780, and by the University of Colorado NeuroTechnology Center. We thank Taylor Yamauchi and other members of the Felsen lab for their comments on the manuscript and Nathan D. Baker for technical assistance. Light microscopy was performed at the University of Colorado Anschutz Medical Campus Advance Light Microscopy Core and engineering support was provided by the University of Colorado Optogenetics and Neural Engineering Core.

